# Positive allosteric modulator of SERCA pump NDC-1171 attenuates cardiac functional decline in mouse model of Duchenne muscular dystrophy

**DOI:** 10.64898/2026.03.05.709950

**Authors:** Niharika Narra, Alyssa M. Richards, Conner C. Earl, Abigail D. Cox, Russell Dahl, Wendy A. Koss, Craig J. Goergen

**Affiliations:** Weldon School of Biomedical Engineering, Purdue University, West Lafayette, Indiana, USA; Indiana University School of Medicine, Indianapolis, Indiana, USA; Department of Comparative Pathobiology, College of Veterinary Medicine, Purdue University, West Lafayette, Indiana, USA; Animal Behavior Core, Purdue University, West Lafayette, Indiana, USA; Neurodon Corporation., Crown Point, Indiana, USA

**Keywords:** Duchenne muscular dystrophy, SERCA activation, cardiac function, pharmacokinetics, D2.*mdx* mouse model

## Abstract

Progressive cardiomyopathy is the leading cause of death in Duchenne muscular dystrophy (DMD). Dysregulation of calcium handling has been implicated in cardiomyopathy progression in DMD. Here we describe a therapeutic approach to improve calcium homeostasis in a mouse model of DMD using the novel therapeutic NDC-1171, which is a positive allosteric modulator of the sarcoplasmic/endoplasmic reticulum calcium ATPase (SERCA) pump. We synthesized NDC-1171 and treated 4-week-old D2.*mdx* mice (n=9) via oral gavage. A group of D2.*mdx* mice (n=9) and a group of DBA/2J mice (n=9; background strain) received a vehicle on the same schedule. We used ultrasound to assess left ventricular function, followed by a treadmill exhaustion test and a 4-paw grip strength test to assess skeletal muscle function. NDC-1171 attenuated cardiac functional decline in D2.*mdx* mice. At 16 weeks of age, left ventricular ejection fraction (LVEF) was significantly preserved in mice treated with NDC-1171 (57.7□±□0.5%) compared to mice treated with a vehicle (50.7□±□0.9%, *p*□<□0.05), though remained lower than background strain controls (62.4□±□0.6%). In contrast, functional behavior testing revealed no significant improvement in skeletal muscle function with treatment. These data suggest that treatment with the SERCA pump modulator NDC-1171 helps preserve cardiac function in a murine model of DMD, even as skeletal muscle function was impaired. Future work will be needed to determine if the benefits of this novel SERCA activator translate to large animal and clinical studies, but these initial results are promising and could help guide development of future treatments for pediatric patients with muscular dystrophy.

## Introduction

Cardiomyopathy is the leading cause of death in Duchenne muscular dystrophy (DMD), an X-linked genetic disorder that affects 1 in 5000 live male births (1). A central driver of DMD-associated cardiac and skeletal muscle dysfunction is impaired calcium handling, including reduced sarcoplasmic/endoplasmic reticulum Ca^2^□ reuptake. The SERCA pump, which normally restores cytosolic Ca^2^□ to the SR during diastole, becomes functionally compromised in dystrophic muscle (2). Enhancing SERCA activity therefore represents a mechanistically targeted approach to improve Ca^2^□ cycling, preserve contractile performance, and counter the progression of cardiomyopathy in DMD (Figure 1).

**Figure 1.**
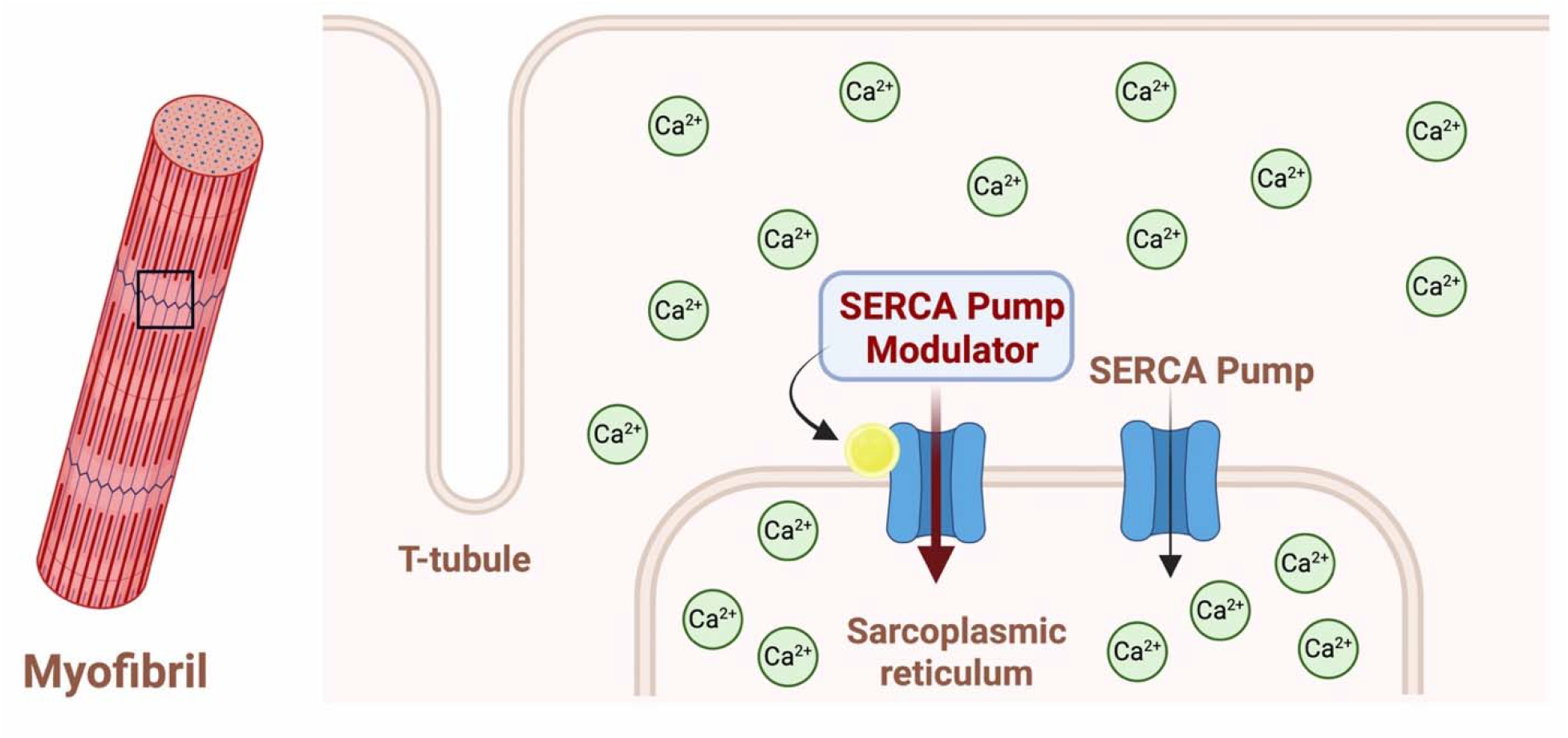
A) Overview of Ca^2^□ cycling in a myofibril, highlighting the relationship between the TLtubule and sarcoplasmic reticulum (SR). During normal excitation–contraction coupling, Ca^2^□ released into the cytosol is cleared primarily through SERCA□mediated reuptake into the SR to support myocyte relaxation. B) Proposed mechanism of action for CDN-1163 and NDC□1171 as a positive allosteric SERCA modulator. A SERCA pump modulator enhances SERCA activity, increasing the rate and efficiency of SR Ca^2^□ reuptake. This augmented Ca^2^□ handling is expected to improve relaxation and partially restore homeostasis in dystrophic muscle where SERCA function is impaired.

The series of quinoline amides that gave rise to CDN-1163 (and later NDC-1171) was discovered via a high-throughput fluorescence resonance energy transfer (FRET) assay consisting of labeled SERCA and phospholamban (3). This assay proved to be more fruitful than previous attempts at finding SERCA activators because it enabled the identification of compounds that caused subtle perturbations in SERCA conformation rather than relying on traditional ATPase-based assays. Medicinal chemistry optimization of this series to enhance its drug properties led to the development of CDN-1163, a molecule that has been the subject of multiple reports in various cellular and animal models revealing that it attenuates neurodegenerative and metabolic disorders (4–7). It is also notable that this series was vetted for off target activity in an MDS Panlabs assay consisting of more than 160 targets (enzymes, receptors, ion channels), including ATPases, and was remarkably selective for SERCA (8). Although the novel SERCA activator CDN-1163 showed reasonable pharmacokinetic parameters in mice (Table 1), the compound needed additional optimization to increase systemic drug levels (AUC, C_max_), reduce clearance (Cl), and boost oral bioavailability (F).

**Table 1.** Pharmacokinetic parameters of CDN-1163 and NDC-1171 after oral dosing in mice (10 mg/kg).

| Compound ID | AUC | $t_{1/2}$ | Cl | $C_{\max}$ | F |
| --- | --- | --- | --- | --- | --- |
|  | (ng•h/mL) | (h) | (mL/min/kg) | (μM) | (%) |
| **CDN-1163** | 283 | 1.17 | 251 | 0.3 | 13.2 |
| **NDC-1171** | 583 | 1.14 | 89 | 1.7 | 30.8 |

**Table 2.**
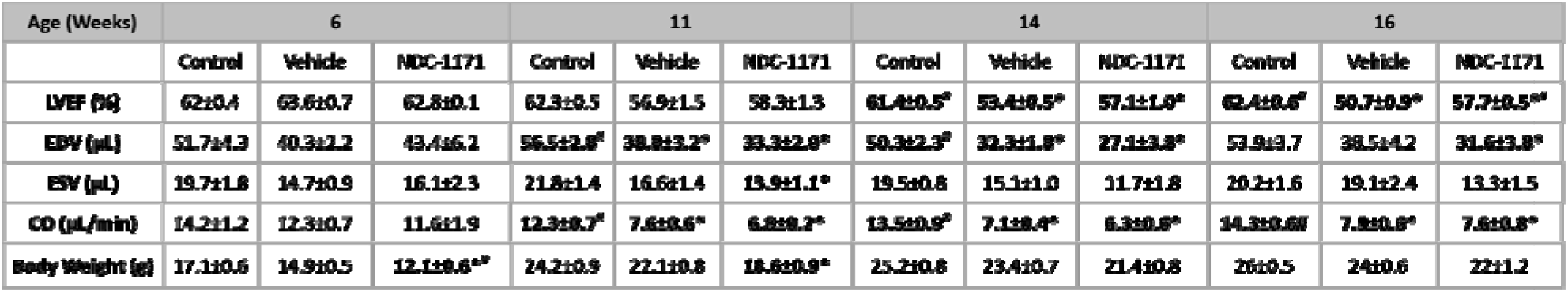
Cardiac function in control, vehicle, and NDC-1171 groups using 2D Ultrasound. Data shown as mean ± SEM. LVEF, left ventricular ejection fraction; EDV, left ventricular end-diastolic volume; ESV, left ventricular end-systolic volume; CO, cardiac output. \**p*<0.05 NDC-1171 different from Control, #*p*<0.05 NDC-1171 differs from Vehicle.

In this study, we aimed to design additional analogues that preserved SERCA activation while meeting key pharmacokinetic properties for effective administration. Our efforts resulted in NDC-1171 which had improved pharmacokinetic properties that met most of our selection criteria and proved to activate SERCA to a greater degree than CDN-1163. With noninvasive echocardiography, treadmill running to exhaustion, and grip strength testing, we were able to characterize the effects of this compound on both cardiac and skeletal muscle. The results suggest that NDC-1171 is a promising and orally available candidate with potential for improved *in vivo* efficacy over CDN-1163 for the treatment of Duchenne-associated cardiomyopathy.

## Methods

### Synthesis of NDC-1171

We initially set out to synthesize additional analogues of CDN-1163 that maintained SERCA activation efficacy while adhering to the following ideal pharmacokinetic criteria: oral bioavailability > 30%, clearance < 100 ml/min/kg, t_1/2_ ≥ 1 hour, and plasma C_max_ > 1µM. Similar to the previously reported synthesis methods of CDN-1163, NDC□1171 was synthesized by first activating a substituted benzoic acid to its acid chloride and then coupling it with 2□methylquinolin□8□amine to form the benzamide scaffold. We purified the crude product by standard silica□based chromatography to remove residual starting materials and minor by□products. We then confirmed compound identity and purity using LCMS and 1H NMR prior to use in downstream assays. Complete experimental details, reaction conditions, and full analytical characterization are provided in the supplementary methods (Appendix A.)

### Ca-ATPase assay

We assessed the effects of CDN-1163 and NDC-1171 on Ca-ATPase activity by measuring the ATP hydrolysis rate using an NADH-linked, enzyme-coupled ATPase assay in 96-well microplates. Each well contained cardiac sarcoplasmic reticulum (SR) membrane vesicles obtained from pig hearts (7 µg), 3-(N-morpholino) propanesulfonic acid (MOPS) (50 mM), MgCl2 (5 mM), KCl (100 mM), NADH (0.2 mM), EGTA (1 mM), phosphoenol pyruvate (1 mM), lactate (5 IU), pyruvate kinase (5 IU), the calcium ionophore A23187 (3.5 µg/mL), and a range of concentrations (1, 3, and 10 µM) for each compound. We added CaCl□ to ensure a saturating Ca^2^□ concentration. We then determined V_max_ by fitting ATPase activity data to the Hill function (9). We initiated the assay by adding ATP to reach a final concentration of 5 mM per well, and measurements were taken using a SpectraMax Plus microplate spectrophotometer.

### Pharmacokinetics

All work with animals described in this manuscript was approved by the Purdue Institutional Animal Care and Use Committee (protocol 1909001948). We assessed the pharmacokinetics of test compounds in male CD-1 mice (n=3 per group) obtained from the Jackson Laboratory. The compounds were formulated at 1 mg/mL in DMSO/Tween 80/water (10/10/80, vol/vol/vol) and dosed at 2 mg/kg (IV) and 10 mg/kg (PO) in triplicate. Blood samples (40 µL) were drawn at 0.25, 0.5, 1, 2, 4, 8, and 24 hours into EDTA containing tubes and plasma was harvested by centrifugation. We added plasma (10 μL) containing 50% acetonitrile (ACN) in water (5 μL) to 200 µL of ACN containing an internal standard. We then vortexed the samples for 30 s. After centrifugation at 4º C and 4,000 rpm for 15 min, the supernatant was diluted 3 times with water. Then, 20 µL of diluted supernatant was injected into the LC/MS system for quantitative analysis. We injected the samples onto an Agilent ZORBAX XDB-Phenyl 5 µm column (50 × 2.10 mm). Mobile Phase A was water with 0.1% trifluoracetic acid (TFA). Mobile Phase B was ACN with 0.1% TFA. Separation was achieved using a gradient of 70% A/30% B to 0% A/100% B over 5.20 min. An AB Sciex LC-MS/MS-EP Triple Quad 6500+ equipped with an electrospray ion spray source was used for all analytical measurements. We measured peak areas of the product ion against the peak areas of the internal standard for quantification, followed by fitting the data using PK Solutions (Summit Research, Montrose, CO).

### DMD Animal model

Male D2.*mdx* mice (n=18, strain #013141, D2.B10-Dmdmdx/J) and its background strain DBA/2J mice (Control) were obtained from Jackson Laboratory at 4 weeks of age (n=9). Mice were group housed and maintained on a 12/12 h light/dark cycle with ad libitum access to food and water. Baseline ultrasound imaging occurred in a subset of mice at 6 weeks of age followed by 8 weeks of drug administration from 6 to 14 weeks of age. During the 4^th^ week of drug administration, at the end of drug administration, and 2 weeks later after behavior tasks, we obtained cardiac ultrasound imaging. Behavior analysis was performed 1 week after the end of drug administration (Figure 2).

**Figure 2.**
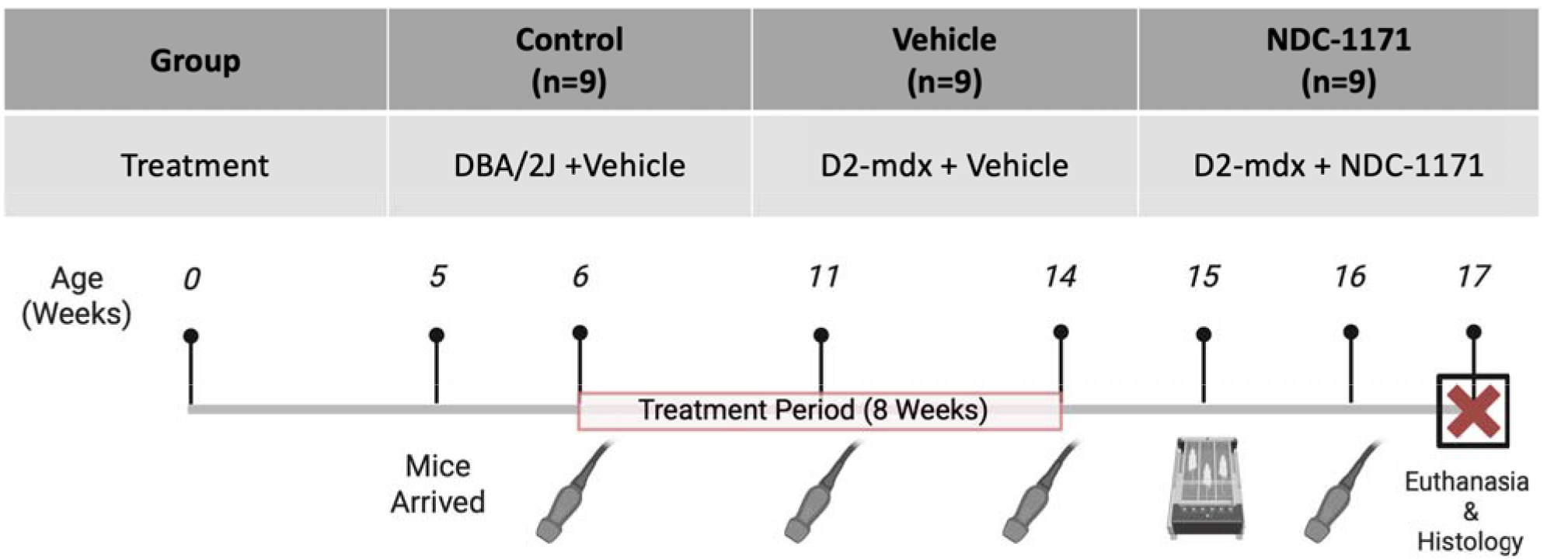
Study design outlining treatment of DBA/2J (control), D2.*mdx* vehicle, and D2.*mdx* NDC□1171 groups (n = 9 per group). Mice received oral gavage from 6 to 14 weeks of age. Ultrasound transducer icons denote timepoints at which transthoracic echocardiography wa performed (baseline: 6, midpoint: 11, endpoint: 14, and post-behavior: 16 weeks of age; n□=□5–6 per group). Behavioral testing icon represents the functional behavior testing (treadmill exhaustion and four□limb grip strength) conducted at week 15. Mice were euthanized at week 17 and cardiac tissue was collected for histological evaluation.

### Drug administration

D2.*mdx* mice were administered vehicle (n=9) or NDC-1171 (n=9) via oral gavage 3 times per week for 8 weeks at a dose of 40 mg/kg in a vehicle containing 10% DMSO/10% Tween80 in water at a 3 mg/mL concentration. Based on the pharmacokinetic results, we selected a treatment dose of 40□mg/kg to achieve the systemic exposures predicted to support sustained SERCA engagement during the study period. These data indicated that higher plasma concentrations were required for effective target coverage over each dosing interval, meaning we adjusted the *in vivo* dosing regimen accordingly. Mice of the background strain, DBA/2J (Control), received vehicle for use as a negative control (n=9). Using this regimen, we observed no adverse effects or weight loss throughout the entire 8 weeks of administration.

### Ultrasound imaging

We anesthetized a subset of mice with 2% isoflurane in an initial induction chamber (Control: n=6; Vehicle: n=5; NDC-1171: n=5). Once each individual animal was anesthetized, it was transferred to an imaging stage and placed in the supine position. We delivered isoflurane through a nose cone to a lowered level of 1.5% over the course of the imaging procedure. The mouse was warmed via the heated stage to a temperature near 36±2 ºC. Body temperature was monitored with a rectal probe. Respiratory rate and heart rate were monitored by the stage which measures inductance changes. Respiratory rate and internal temperature were monitored to ensure proper anesthesia and heating was maintained.

We performed transthoracic echocardiography at the baseline, midpoint, endpoint, and post-behavior testing timepoints (Figure 2). We used a Vevo 3100 small animal ultrasound system (FUJIFILM VisualSonics Inc.) with the MX550D linear array transducer (25-55 MHz). We removed thoracic hair with depilatory cream prior to imaging. We obtained long- and short-axis EKG-gated kilohertz visualization and long and short axis motion–mode (M-mode) images. From these images, we calculated left ventricular ejection fraction as a % (LVEF), left ventricular end diastolic and left ventricular end systolic volume in µL (LVEDV and LVESV, respectively), and cardiac output in µL/min (CO).

### Functional Behavior Testing

#### Treadmill exhaustion test

Mice were tested on a 5-lane treadmill (Panlab LE8710MTS) with a 10-degree upward angle. They were trained for 3 days for 5 minutes at a speed of 6 m/min. During testing mice began at a speed of 6 m/min which gradually increased by 1.2 m/min every minute until exhaustion. Exhaustion was defined as the time when the mouse could no longer run despite repeated mechanical stimulation with soft paddles (4 cm^2^) that were incorporated into each lane of the treadmill.

#### 4-paw grip strength

The combined grip of the forelimbs and hindlimbs was recorded using a grip strength meter. The grip strength meter was positioned horizontally, and the mouse was held by its tail and allowed to securely grip the metal grid with all 4 paws. After the mouse obtained a solid grasp of the metal grid, the mouse was pulled backward by the base of its tail onto the experimenter’s forearm. The force that was applied to the grip strength meter at the time of release was recorded as the peak grip strength. The measurement was repeated 5 times, and the average force was determined for each mouse. Grip strength values (grams of force) were normalized to the body weight (grams) of each mouse (g/g BW).

### Histology

At the conclusion of the study period, each mouse was anesthetized and maintained on 4% isoflurane. Once anesthetized, we surgically opened the chest and injected 1 M KCl solution into the apex of the heart arresting it in diastole. We then excised the hearts and perfused briefly with cold PBS, followed by 4% paraformaldehyde. Each heart was then sliced at the mid papillary level in the short axis, embedded in paraffin and stained with hematoxylin and eosin (H&E) and Masson’s trichrome. We imaged slides with stained cross sections at 4x magnification. A veterinary pathologist evaluated the slides for fibrosis and necrosis.

### Statistical Analysis

Statistical analyses were performed using GraphPad Prism 10.6.1 (GraphPad Software Inc., San Diego, CA, US). Pharmacokinetic parameters were determined using non□compartmental analysis with the PK Solutions 2.0.2 software package (SUMMIT Research Service, Montrose, CO). Echocardiography data were analyzed using a two-way ANOVA with Tukey’s multiple comparisons test. Functional behavior outcomes from grip strength and treadmill exhaustion tests were both analyzed using a one□way ANOVA with Dunnett’s multiple comparisons test, with Vehicle as the comparator. We evaluated histological data qualitatively and did not perform any statistical analysis

## Results

### Pharmacokinetics

In cardiac SR vesicles, both CDN-1163 and NDC-1171 increased SERCA V_max_ in a dose-dependent manner, with NDC-1171 producing larger V_max_ shifts at matched concentrations (Figure 3). *In vivo* pharmacokinetics in mice demonstrated that NDC-1171 achieved higher exposure (AUC 583 ng•h/mL; C_max_ 1.7µM), lower clearance (Cl 89 mL/min/kg), and a greater oral bioavailability (F 30.8%) compared with CDN-1163 (Table 1). Taken together, stronger SERCA activation and a more favorable exposure profile for NDC□1171 provide the mechanistic and PK/PD rationale for prioritizing this analogue in the subsequent *in vivo* efficacy experiments in D2.*mdx* mice.

**Figure 3.**
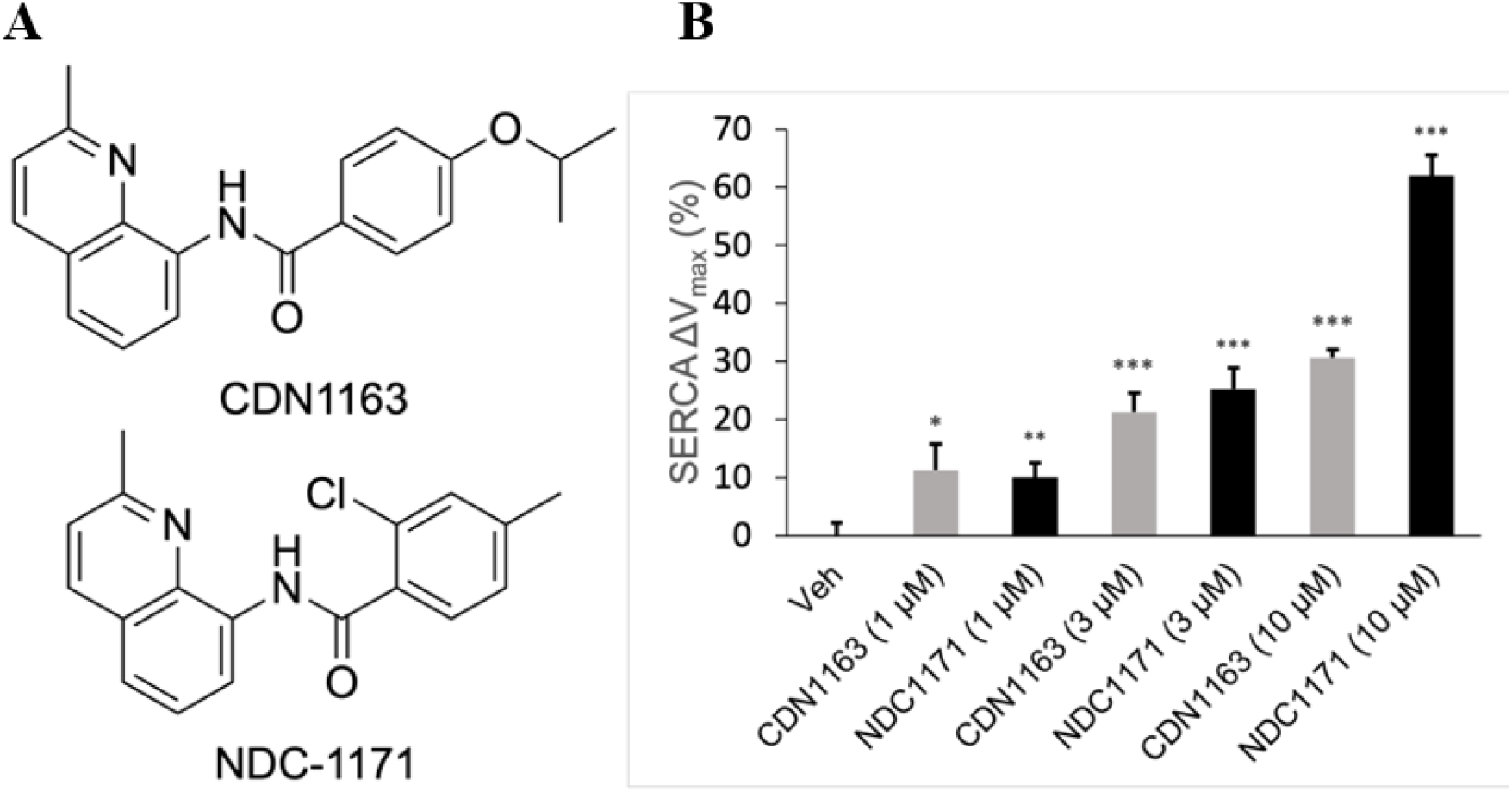
(A) Structures of CDN-1163 and NDC-1171. (B) Effects of increasing doses of CDN-1163 and NDC-1171 on SERCA V_max_ in cardiac SR vesicles. Mean ± S.E. (n = 6). \**p* < 0.05; \*\**p* < 0.01; \*\*\**p* < 0.001.

### Cardiac function

At baseline, all groups had the same global ventricular function, with similar ejection fraction, end diastolic volume, and end systolic volume. At 14 weeks, the untreated D2.*mdx* group’s ejection fraction declined significantly from that of the DBA/2J. At 16 weeks, all three groups were significantly different (*p*<0.001; Figure 4) with LVEF control = 62.4 ± 0.6%, vehicle = 50.7 ± 0.9%, and NDC-1171 = 57.7 ± 0.5%.

**Figure 4.**
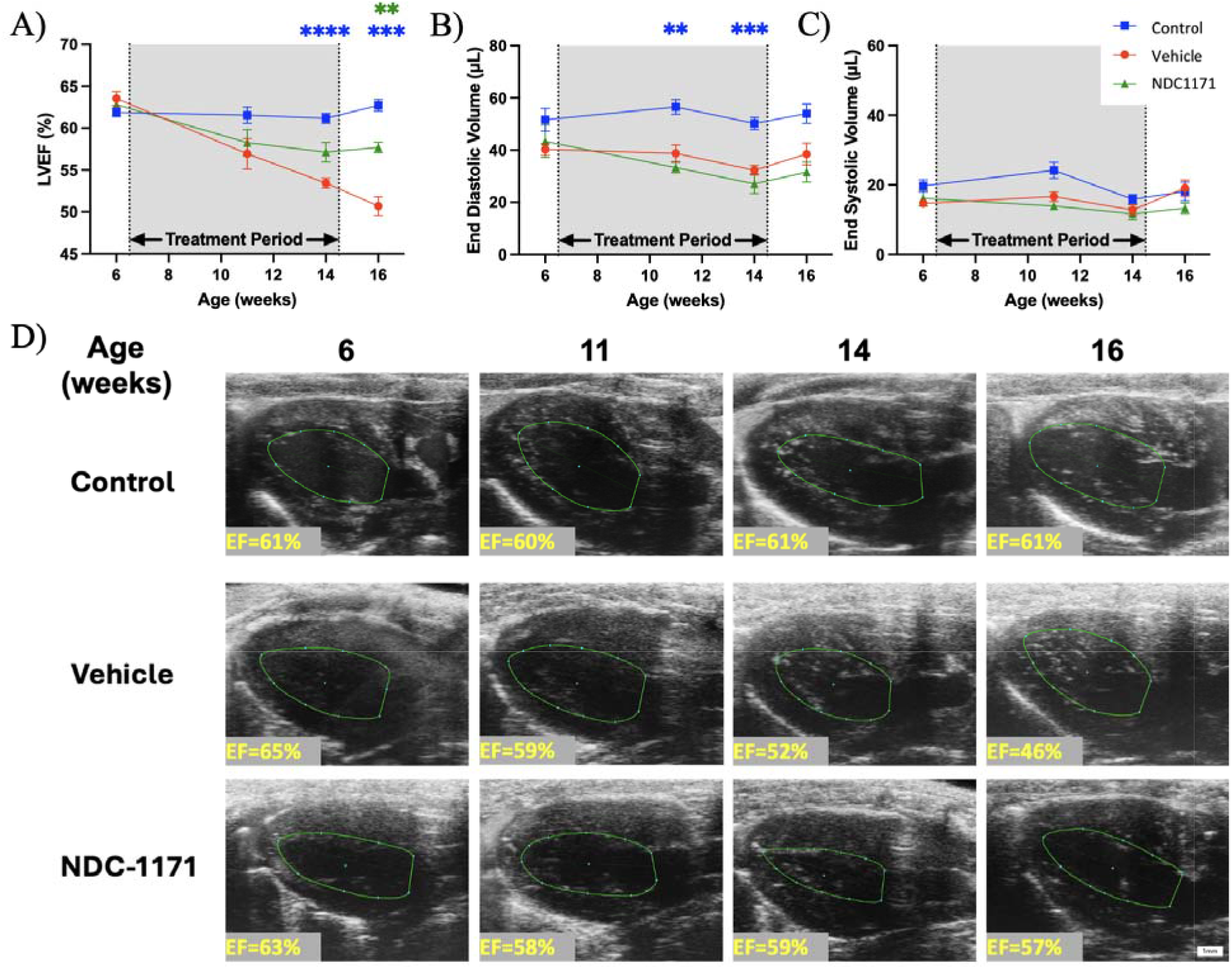
NDC-1171 attenuates cardiac functional decline in a *mdx* mouse model. A) left ventricular ejection fraction, B) end diastolic volume, C) end systolic volume tracked longitudinally in Control (blue), Vehicle (red), and NDC□1171 (green) groups. Data shown as Mean ± S.E. where \*\**p*<0.01 and \*\*\**p*<0.001 (Blue = Control significantly different from Vehicle; Green = NDC□1171 significantly different from Vehicle). D) A representative long axis image from each group at end diastole. Scale bar: 1 mm.

### Behavioral Studies

NDC-1171 did not attenuate the decrease in muscle strength and running stamina in D2.*mdx* mice as shown by 4-limb grip strength test and the treadmill exhaustion test (Figure 5). There were no significant differences between groups treated and untreated groups on either the grip strength (*p*=0.90) or treadmill (*p*=0.64) test.

**Figure 5.**
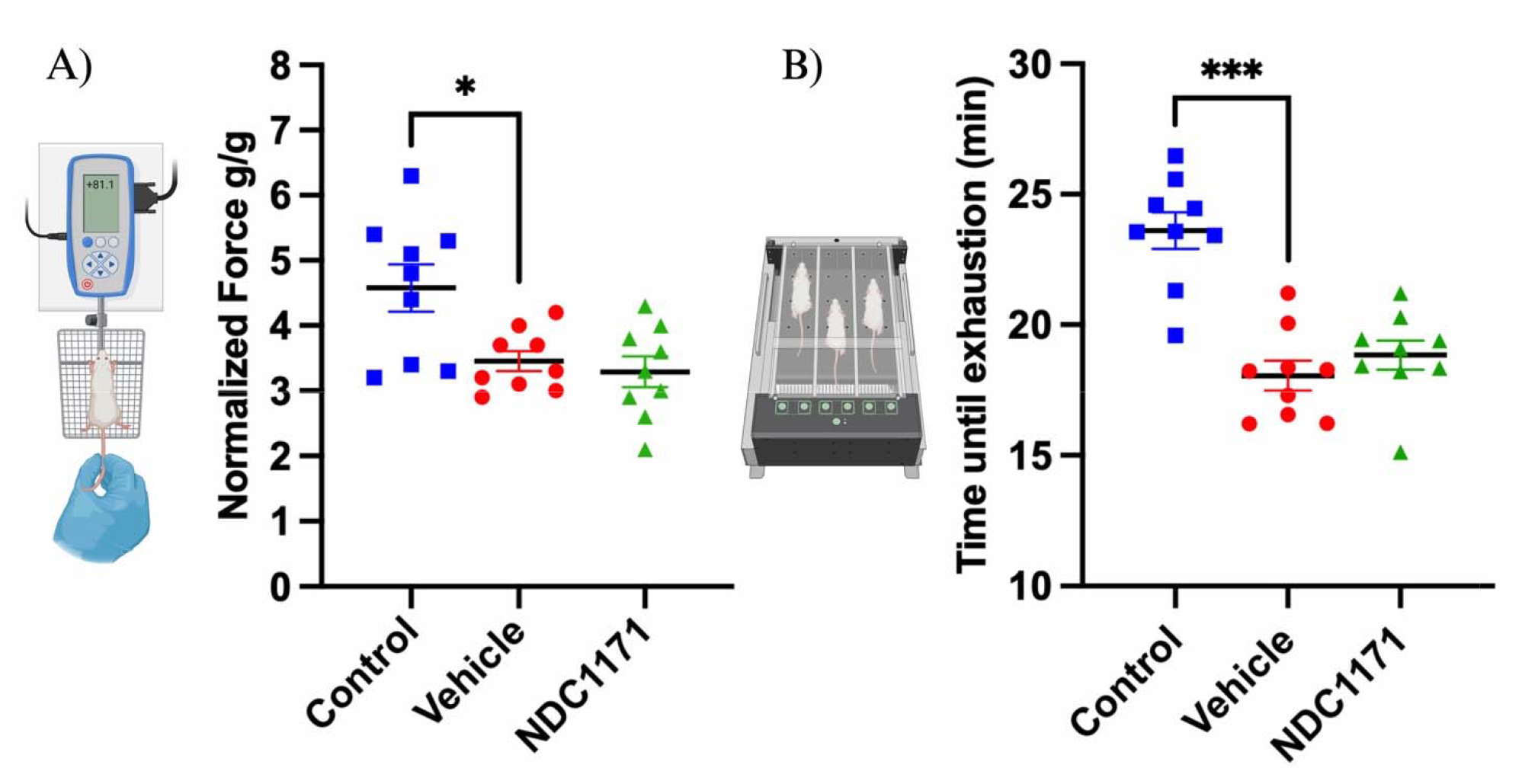
D2.*mdx* vehicle-treated mice (Vehicle) had impairments when compared to DBA/2J vehicle-treated (Control) in both (A) Four-limb grip strength test and (B) Treadmill exhaustion test. D2.*mdx mice treated with* NDC-1171 did not improve in either behavior assessments. \**p*<0.05, \*\**p*<0.01, and \*\*\**p*<0.001.

**Figure 6.**
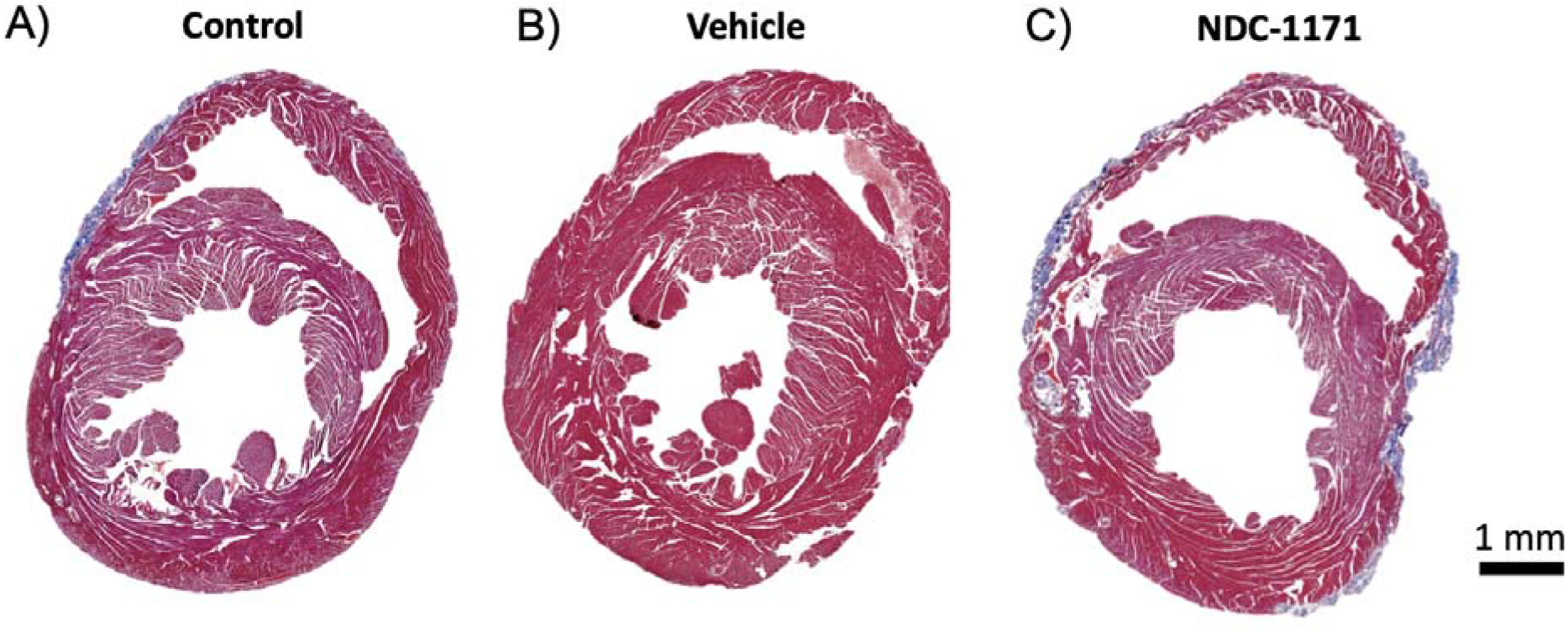
A representative mid-papillary coronal section of A) Control, B) Vehicle, and C) NDC-1171 left ventricles stained with Masson’s trichrome show no significant myocardial fibrosis in any group. Scale bar: 1 mm.

### Histology

There was no conspicuous presence of fibrosis seen in any of the samples. A fibrotic response was identified histologically along the interface of pericardium/epicardium and myocardium. This pericardial fibrosis was identified in control mice as well.

## Discussion

In this study, we sought to determine the potential benefit of NDC-1171, a novel SERCA activator (SERCA), in supporting cardiac and behavioral decline in a murine model of DMD. Our results show a potential cardiac functional benefit due to this treatment, likely mediated by SR cycling. The ejection fraction of treated mice was significantly higher than that of the untreated vehicle group at week 16 (Figure 4). This observed preservation of systolic function is consistent with expected effects of enhanced sarcoplasmic/endoplasmic reticulum Ca^2+^ reuptake via SERCA2a, which is the predominant isoform in the cardiac muscle. This lowers cytosolic Ca^2+^ during diastole and helps maintain Ca^2+^ load for subsequent cardiac contraction. Duchenne-associated cardiomyopathy is characterized by Ca^2+^ mishandling across the sarcoplasmic/endoplasmic reticulum and mitochondria (10). Therefore, a pharmacologic approach that restores Ca^2+^ cycling could address a core pathophysiologic mechanism (11).

Interestingly, histological analysis did not show differences in fibrosis between groups. This suggests that the functional benefit of NDC-1171 in *mdx* mice is not due to the prevention of fibrotic replacement of the myocardium. While fibrosis was not significantly different, this may reflect sampling variability or the relatively short duration of treatment. Quantitative collagen metrics and longer study duration could clarify whether SERCA activation influences fibrosis and cardiac remodeling. These observations should also be interpreted in the context of the D2.*mdx* cardiac phenotype, where chamber-specific remodeling, particularly in the epicardium of the right ventricle along with significant inter-animal variability have been reported (12,13). Thus, additional work is needed to elucidate the relationship between of cardiac fibrosis function in preclinical muscular dystrophy studies.

We did not observe any attenuation in skeletal muscle function decline in the treated group. The treadmill test results do not provide any evidence that the cardiac reserve in treated mice improves the exercise tolerance in this group, nor does the grip strength test suggest any preservation of skeletal muscle function. This suggests that NDC-1171 treatment inotropically supports cardiac function without preventing remodeling or impacting the skeletal muscle function. The preserved ejection fraction, despite the lack of skeletal muscle improvement or cardiac muscle remodeling, suggests tissue-specific pharmacokinetics or isoform differences in cardiac and skeletal muscle SERCA. These findings warrant future studies on drug distribution and target engagement.

Previously, it has been shown that the SERCA pump activator CDN-1163 can ameliorate dystrophic phenotypes in a murine *mdx* model (14). This includes stronger skeletal muscles, reduced degeneration and fibrosis, and protection against exercise-induced damage after treatment with CDN-1163 for 3 weeks. Those findings demonstrated that SERCA activation with CDN-1163 could be a therapeutic strategy for the skeletal muscle symptoms of DMD. As the next-generation analogue, NDC-1171 was developed to preserve SERCA-activating efficacy while improving pharmacokinetic properties. It would therefore be hypothesized that NDC-1171 should then also have a therapeutic effect on skeletal muscle. Our data show preserved cardiac ejection fraction with NDC-1171 without any measurable skeletal muscle improvement, suggesting that SERCA activation can yield organ-specific windows of efficacy. Differences in pharmacokinetics, including higher C_max_ and bioavailability of NDC-1171 compared to CDN-1163 (Table 1), may contribute to these distinct response profiles. A plausible explanation is the upregulation of sarcolipin (SLN) in dystrophic skeletal muscle, which interferes with the ability of SERCA to couple Ca^2+^ transport with ATP hydrolysis. Additionally, prior efficacy with CDN□1163 was reported following intraperitoneal administration, whereas our study used oral dosing with NDC□1171 to reflect intended clinical use. Differences in absorption and first□pass metabolism between routes may have contributed to the different response profiles across cardiac versus skeletal muscle. This uncoupling reduces net Ca^2+^ uptake even in the presence of a SERCA activation, whereas cardiac SERCA2a is primarily regulated by phospholamban (15,16). Future work, while outside of the scope of this initial study, could include dose response studies that investigate optimal treatment period duration and dosages.

It is also possible that the exercise studies exacerbated the cardiac decline between the endpoint and post-behavior timepoints in this study. The decline between endpoint and post-behavior was only pronounced in the vehicle group, which suggests that the NDC-1171 treatment may also preserve function in the context of cardiac stressors. We also note that while previous studies have also shown that D2.*mdx* mice have an increase in epicardial cardiac fibrosis compared to DBA controls at around 16-18 weeks of age (12,13), we did not detect significant fibrosis in either the treated or untreated D2.*mdx* groups. Variations between D2.*mdx* mice have been shown to be large, with some animals having almost no fibrosis and others having pronounced fibrosis (17). Additional physical or pharmacological stressors could be used to exacerbate cardiomyopathy and fibrosis in the *mdx* mouse model. For instance, both physical exercise (18) and low□dose β□adrenergic stimulation with isoproterenol have been used to reveal reduced systolic responsiveness and stress□induced dysfunction in *mdx* mice (19), suggesting cardiac stress plays a role in disease progression and cardiac remodeling.

This study has important limitations that should be considered when interpreting the findings. First, the sample size for functional echocardiographic assessment was relatively small, which may reduce the statistical power to identify differences across groups. Additionally, diastolic function was not assessed in this study; incorporating diastolic measurements may help determine whether SERCA activation improves early filling dynamics. Nevertheless, we still observed significantly better cardiac function in the treatment group compared to vehicle□treated controls. Second, histological evaluation was based on single mid□left ventricular cross□sections, which may miss regional remodeling previously reported in D2.*mdx* mice. Additional techniques for quantifying cardiac fibrosis in tissue *ex vivo* may help address this limitation. Finally, the eight□week study duration may have been insufficient to reveal structural, molecular, or behavioral benefits that develop more slowly. The D2.*mdx* strain also exhibits an earlier and more severe cardiac phenotype compared to other murine models and what is typically observed in many patients, potentially influencing our ability to discern treatment effects. Future studies could consider alternative DMD models such as the B10□*mdx* (C57BL/10□*mdx*) strain to better reflect baseline severity and the slower progression of the cardiac phenotype (19,20). Expanded cohort sizes, extended dosing regimens, and physiologic stress testing will also be important for developing a more complete therapeutic profile of NDC□1171. Longer study durations, quantitative fibrosis assessments, and incorporation of diastolic function measurements may help clarify additional structural or functional effects that were not detectable in this study. These follow□up investigations will be important for determining how sustained exposure, optimized dosing, or alternative disease models may more comprehensively evaluate the therapeutic potential of NDC□1171 in Duchenne□associated cardiomyopathy.

## Conclusion

Overall, these findings demonstrate that NDC-1171 provides measurable cardio protection in the D2.*mdx* model of DMD by preserving left-ventricular systolic function during a critical window of disease progression in dystrophic hearts. The observed cardiac benefit, in the absence of measurable myocardial fibrosis, suggests that NDC-1171 exerts its effect primarily through the modulation of calcium dynamics rather than structural remodeling in the early disease stages. By improving SERCA□mediated Ca^2^□ cycling and exhibiting pharmacokinetic properties superior to earlier compounds in this series, NDC□1171 represents a promising next□generation therapeutic candidate for Duchenne□associated cardiomyopathy. The cardiac□dominant pattern of efficacy also underscores the need for extended treatment durations, expanded cohort sizes, and comprehensive functional and biochemical analyses to determine whether broader benefits, particularly in skeletal muscle, emerge with sustained exposure or optimized dosing for DMD patients.

## Appendix A

### Synthesis of CD-1163

Oxalyl chloride (6.0 mL, 2M in DCM) was added dropwise into a stirred solution of 4-isopropoxybenzoic acid (1.80g, 9.989 mmol) and 50.0 µL of dimethylformamide (DMF) in DCM (30.0 mL) at 0□ under nitrogen. The resulting mixture was stirred for 1 hour at room temperature and then concentrated under reduced pressure. The crude product was used in the next step directly without further purification. The residue was dissolved in DCM (30.0 mL) and added dropwise into a stirred solution of 2-methylquinolin-8-amine (1.70 g, 10.746 mmol) and TEA (2.17 g, 21.445 mmol) in DCM (30.0 mL) at 0 □. The resulting mixture was stirred for 2 hours at room temperature under a nitrogen atmosphere. Liquid chromatography – mass spectrometry (LCMS) showed the starting material was consumed and the product was formed. The resulting mixture was washed with water and dried over anhydrous Na2SO4. After filtration, the filtrate was concentrated under reduced pressure. The residue was purified using silica gel column chromatography eluting with EtOAc/DCM (0-5%) to afford crude product. The crude product was dissolved in DCM (50 mL), washed with 1N HCl (50 ml), and washed with 10% NaHCO3 (50 ml). The organic layers were dried over anhydrous Na2SO4. After filtration, the filtrate was concentrated under reduced pressure to afford 1.8 g (52% yield) of the title compound as a white solid. LC-MS: (ESI, m/z): [M+H] + = 321. 1H NMR (400 MHz, DMSO-d6) δ 10.63 (s, 1H), 8.71 – 8.67 (m, 1H), 8.33 (d, J = 8.4 Hz, 1H), 8.01 – 7.95 (m, 2H), 7.68 – 7.63 (m, 1H), 7.59 – 7.52 (m, 2H), 7.13 (d, J = 9.2 Hz, 2H), 4.82 – 4.72 (m, 1H), 2.78 (s, 3H), 1.32 (d, J = 6.0 Hz, 6H).

### Synthesis of NDC-1171

NDC-1171 was synthesized by adding oxalyl chloride (2M in DCM) (3.9 mL, 7.800 mmol) to a solution of 2-chloro-4-methylbenzoic acid (903.1 mg, 5.294 mmol) in DCM (10 mL) along with DMF (35.9 mg, 0.491 mmol) at 0 ºC. The resulting mixture was then stirred for 1 h at room temperature under nitrogen. This was then concentrated under vacuum to provide 2-chloro-4-methylbenzoyl chloride. The 2-chloro-4-methylbenzoyl chloride was redissolved in DCM (5 mL) and added dropwise to a solution of 2-methylquinolin-8-amine (800.0 mg, 5.057 mmol) and triethylamine (1.1 g, 10.871 mmol) in DCM (10 mL) at 0 ºC. The resulting mixture was subsequently stirred for 1 h at room temperature under nitrogen until the reaction was shown to be complete by LCMS. The resulting mixture was quenched with 50 mL of water and extracted with DCM (30mL). The organic layer was dried over anhydrous sodium sulfate and concentrated. The residue was applied onto a silica gel column and eluted with ethyl acetate/petroleum ether (1:3) to afford 1.1925 g (77%) of the product 2-chloro-4-methyl-N-(2-methylquinolin-8-yl) benzamide as a white solid. LCMS (ESI): [M+H] = 311. 1H NMR (400 MHz, DMSO-d6) δ 10.52 (s, 1H), 8.73 (d, J = 8.0 Hz, 1H), 8.33 (d, J = 8.8 Hz, 1H), 7.77 – 7.69 (m, 2H), 7.60 – 7.53 (m, 2H), 7.49 (s, 1H), 7.36 (d, J = 7.6 Hz, 1H), 2.68 (s, 3H), 2.40 (s, 3H).

## Data Availability

Data presented in this study are available from corresponding author upon reasonable request.

## Acknowledgments

The authors declare that they have no additional acknowledgements.

## Author Contributions

N.N.: Data curation, formal analysis, writing – original draft, writing – review and editing; A.M.R.: Investigation, formal analysis, writing – original draft, writing – review and editing; C.C.E.: Investigation, methodology, writing – review and editing; A.D.C.: Formal analysis, writing – review and editing; R.D.: Conceptualization, investigation, data curation, formal analysis, writing – review and editing; W.A.K.: Conceptualization, investigation, data curation, formal analysis, writing – review and editing; C.J.G.: Conceptualization, investigation, supervision, writing – review and editing.

